# ProSST: Protein Language Modeling with Quantized Structure and Disentangled Attention

**DOI:** 10.1101/2024.04.15.589672

**Authors:** Mingchen Li, Pan Tan, Xinzhu Ma, Bozitao Zhong, Huiqun Yu, Ziyi Zhou, Wanli Ouyang, Bingxin Zhou, Liang Hong, Yang Tan

**Affiliations:** Shanghai Jiao Tong University, China; Shanghai Artificial Intelligence Laboratory, China; East China University of Science and Technology, China

**Author notes:** Corresponding authors. {, }. Equal contribution (Mingchen Li, Yang Tan).

## Abstract

Protein language models (PLMs) have shown remarkable capabilities in various protein function prediction tasks. However, while protein function is intricately tied to structure, most existing PLMs do not incorporate protein structure information. To address this issue, we introduce ProSST, a Transformer-based protein language model that seamlessly integrates both protein sequences and structures. ProSST incorporates a structure quantization module and a Transformer architecture with disentangled attention. The structure quantization module translates a 3D protein structure into a sequence of discrete tokens by first serializing the protein structure into residue-level local structures and then embeds them into dense vector space. These vectors are then quantized into discrete structure tokens by a pre-trained clustering model. These tokens serve as an effective protein structure representation. Furthermore, ProSST explicitly learns the relationship between protein residue token sequences and structure token sequences through the sequence-structure disentangled attention. We pre-train ProSST on millions of protein structures using a masked language model objective, enabling it to learn comprehensive contextual representations of proteins. To evaluate the proposed ProSST, we conduct extensive experiments on the zero-shot mutation effect prediction and several supervised downstream tasks, where ProSST achieves the state-of-the-art performance among all baselines. Our code and pretrained models are publicly available ^2^.

## 1 Introduction

Predicting the functions of proteins is one of the most critical areas in life sciences [1]. In recent decades, protein sequence databases have experienced exponential growth [2], making it possible to learn the fundamental representations of protein sequences with large-scale models in a data-driven manner. Inspired pre-trained language models in natural language processing [3, 4], many pre-trained Protein Language Models (PLMs) have emerged [5, 6, 7, 8, 9]. Benefiting from remarkable protein representation capabilities, they have become fundamental tools for bioinformatics in protein-related tasks.

According to the central dogma [10], the function of a protein is determined by its structure. However, most PLMs mainly focus on modeling protein sequences, neglecting the importance of structural information, and one significant reason for this phenomenon is the lack of structural data. Fortunately, some excellent works, such as AlphaFold [11] and RoseTTAFold [12], are proposed, which can accurately predict protein structures. These works significantly expand the protein structure dataset [13] to millions and enable the pre-training of large-scale structure-aware PLMs. After that, the major challenge is how to effectively integrate protein structure information into PLMs. Specifically, existing structure-aware PLMs [14, 15] first use Foldseek [16] to convert protein structures into discrete structure tokens and then integrate these structural data into the Transformer architecture. However, despite achieving promising performance on several tasks, this approach still faces two main issues. First, Foldseek encodes the structure of a residue within a protein by considering only the features of its previous and next residues. This representation is insufficient and may overlook subtle differences in the local structure of proteins, such as catalytic sites or binding pockets, which are crucial for protein function [17]. Second, the naive Transformer architecture lacks the ability to explicitly model the relationship between protein sequences and structure token sequences, making it challenging to effectively leverage structural cues.

In this paper, we develop ProSST (**Pro**tein **S**equence-**S**tructure **T**ransformer), a structure-aware pre-trained protein language model. Specifically, ProSST mainly consists of two modules: a structure quantization module and a Transformer with sequence-structure disentangled attention. The structure quantization module is based on a GVP (Geometric Vector Perceptron) [18] encoder, which can encode a residue along with its neighborhoods in its local environment and quantize the encoding vectors into discrete tokens. Compared to Foldseek, which only considers individual residues, this encoder can take into account more information from the micro-environment of residue. The sequence-structure disentangled attention module replaces the self-attention module in the Transformer model. This can make Transformer model explicitly model the relationship between protein sequence tokens and structure tokens, enabling it to capture more complex features of protein sequences and structures. To enable ProSST to learn the contextual representation comprehensively, we pre-train our model with the Masked Language Modeling (MLM) objective on a large dataset containing 18.8 million protein structures. To summarize, our main contributions are as follows:

- We propose a protein structure quantizer, which can convert a protein structure into a sequence of discrete tokens. These token sequences effectively represent the local structure information of residues within a protein.
- We propose a disentangled attention mechanism to explicitly learn the relationship between protein structure and residue, facilitating more efficient integration of structural token sequences and amino acid sequences.

To evaluate the proposed ProSST, we conduct extensive experiments on zero-shot mutation effect prediction and multiple supervised downstream tasks, where the proposed model achieves state-of-the-art results among all baselines. Besides, we also provide detailed ablations to demonstrate the effectiveness of each design in ProSST.

## 2 Related Work

### 2.1 Protein Representation Models

Based on the input modality, protein representation models can be divided into three categories: sequence-based models, structure-based models, and structure-sequence hybrid models.

#### Sequence-based models

Sequence-based models treat proteins as a sequence of residue tokens, using the Transformer model [19] for unsupervised pre-training on extensive datasets of sequence. According to the pre-training objective, current models can be further divided into BERT-based models [4], GPT-based models [3], and span-mask based models. Specifically, BERT-style models, including ESM-series models [5, 6, 7], ProteinBert[9], and TAPE [20], aim to recover the masked tokens in the training phase. The GPT-style models, such as Tranception [21], ProGen2 [22], and ProtGPT2 [23], progressively generate the token sequences in an auto-regressive manner. Lastly, models that use span-mask as the training objective include Ankh [24], ProtT5 [8], and xTrimo [25].

#### Structure-based models

Protein structures play a dominant role in protein functionality. Therefore, models leveraging structure information generally get more accurate predictions. Recently, various techniques have been applied in learning protein structure representation, including CNN-based models [26] and GNN-based models [18, 27, 28, 29], and the GNN-based ones have demonstrated significant versatility in integrating protein-specific features through node or edge attributes. More-over, the recent advancements in protein folding models [11, 7, 30] enable the structure-based models access to extensive datasets of protein structures. This led to a growing interest in developing PLMs that leverage protein structure cues [15, 14, 28].

#### Structure-sequence hybrid models

Hybrid models, which incorporate both sequence and structure information of proteins, offer more effective representations of proteins. For example, the LM-GVP[31] model employs ProtBERT-BFD [9] embeddings as input features for the GVP [18] model, while ESM-GearNet [32] investigates various methods of integrating ESM-1b [5] representations with GearNet [28]. Similarly, the recent ProtSSN [33] model leverages ESM-2 [7] embeddings as input for the EGNN [34] model, resulting in notable advancements. Both ESM-IF1 [35] and MIF-ST [36] target inverse folding, utilizing the structure to predict corresponding protein residues, whereas ProstT5 [15] focuses on the transformation between residue sequences and their structure token sequences [16] as a pre-training objective. SaProt [14] constructs a structure-aware vocabulary using structure tokens generated by foldseek [16]. Both SaProt and ProstT5 extensively utilize large structure databases [13] for their pre-training datasets. ProSST is also a hybrid structure-sequence model. Compared to previous work, ProSST develops an advanced structure quantization method and a better attention formulation to leverage the structure cues.

### 2.2 Protein Structure Quantization

The most intuitive way to represent a protein structure is using continuous features, such as coordinates, dihedral angles and distance map. However, directly using these continuous features in the pre-training may lead to overfitting [14]. This issue arises from the mismatched representations of the structure between the training set (derived from model predictions) and the test set (measured by wet-lab experiments). As the bridge to eliminate this gap, structure quantization has been investigated by a few works. These methods can be divided into two groups based on the way to generate the discrete secondary structure, including the methods based on physical computing, such as DSSP [37], and the methods based on deep learning, such as Foldseek [16], which have been successfully applied to structure-aware PLMs [14, 15]. The structure quantization module of ProSST also relies on learning-based approaches but provides a more detailed residue structure representation, compared to Foldseek.

## 3 Method

In this section, we introduce the architecture of ProSST. ProSST mainly contains two modules: structure quantization (Section 3.1) module and a-transformer-based model with sequence-structure disentangled attention. (Section 3.2).

### 3.1 Structure Quantization Module

The structure quantization module aims to transform a residue ‘s local structure in a protein into a discrete token. Initially, the local structure is encoded into a dense vector using a pre-trained structure encoder. Subsequently, a pre-trained k-means clustering model assigns a category label to the local structure based on the encoded vector. Finally, the category label is assigned to the residue as the structure token. The pipeline of structure quantization is shown in Figure 1.

**Figure 1.**
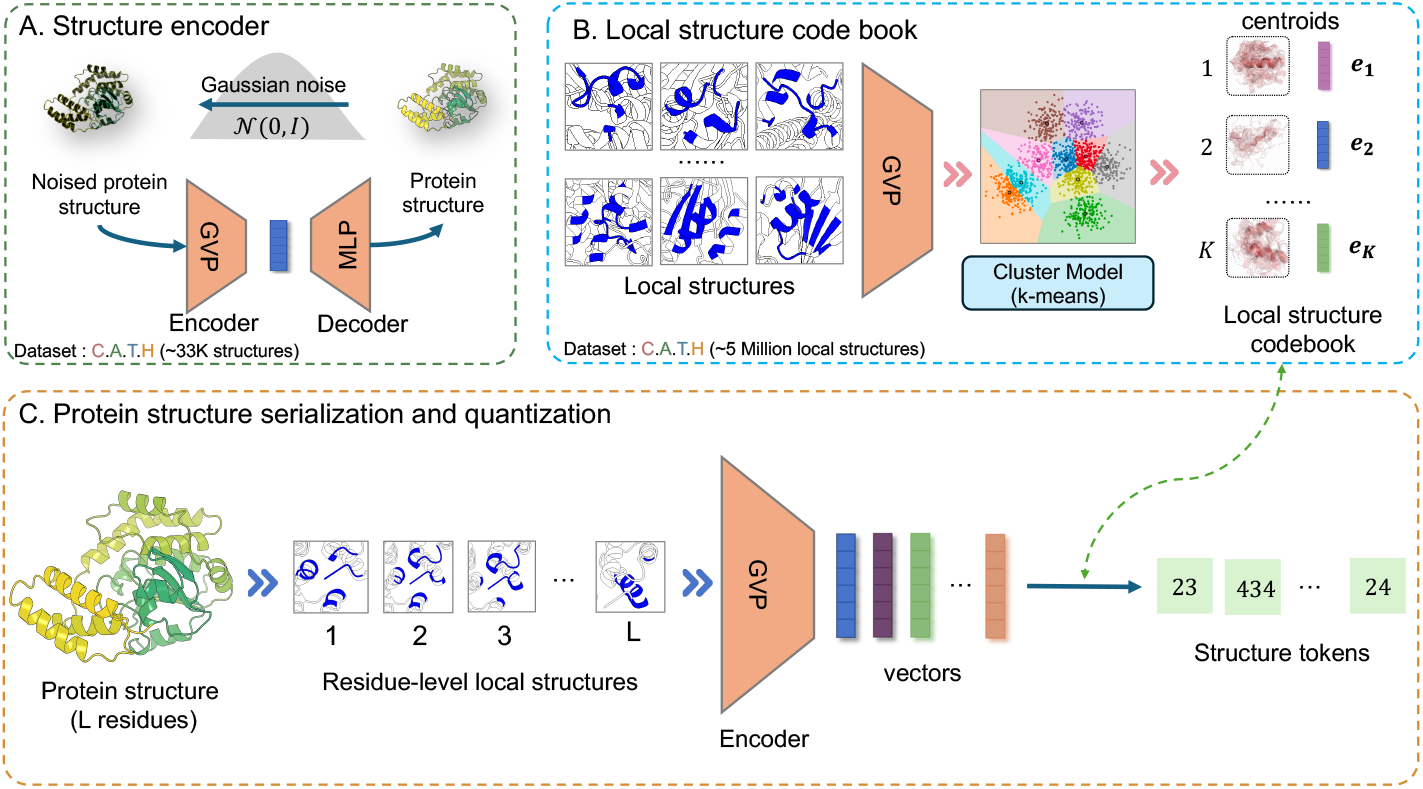
The pipeline of structure quantization. (A) Training of the structure encoder. (B) Local structure clustering and labeling. (C) Converting a protein structure to structure token sequence.

#### Structure representation

We categorize protein structures into two distinct levels: *protein structure* and *local structure*. Protein structure denotes the complete architecture of a protein, encompassing all its residues. The local structure focuses on specific individual residues. It describes the local environment of a residue by centering on a specific residue and including it along with the nearest 40 residues surrounding it in three-dimensional space [18]. Compared to protein structure, local structures are in finer granularity, which allows for a more accurate description of the structure of residue. Therefore, a protein containing *L* residues has one protein structure and *L* local structures. Despite the different levels of structure, we can use graphs to represent it. Formally, we represent a structure using graph ***G*** = (***V***, ***E***), where ***V*** and **E** denote the residue-level nodes and edges, respectively. For any given node ***v*** ∈ ***V***, it contains only the coordinate information of the residue, without any type information of the residue itself. This ensures that the structure encoder is solely focused on the structure cues. The edge set ***E*** = {***e***_*ij*_} includes all *i, j* for which ***v***_*j*_ is one of the forty nearest neighbors of *v*_*i*_, determined by the distance between their ***Cα*** atoms.

#### Structure encoder

Based on the above-mentioned definition of structure, we use geometric vector perceptrons (GVP) [18] as the (local) structure encoder. In particular, the GVP can be represented as a structure feature extraction function *π*_*θ*_(***G***) ∈ ℝ^*l×d*^, where *l* is the number of nodes, *d* is the embedding dimension, and *θ* is trainable parameters. We integrate GVP with a decoder that includes a position-wise multi-layer perceptron (MLP) to form an auto-encoder model. The entire model is trained using a de-noising pre-training objective. In this process, we perturb ***Cα*** coordinates with 3D Gaussian noise (Figure 1A) and use Brownian motion on the manifold of rotation matrices, according to RF-Diffusion [38]. The model is then tasked with recovering the structure to its original, noise-free state. After being trained on the C.A.T.H dataset [39] (see Appendix A.2), we exclude the decoder and utilize solely the mean pooled output of the encoder as the final representation of structures. Although the structure encoder is trained on protein structures, it can effectively be applied to local structures. Therefore, for a specified graph ***G***, the encoding process can be described by: 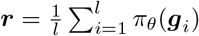, where ***g***_*i*_ represents the features of the local structure associated with the *i*-th node in the graph ***G***, and *π*_*θ*_(***g***_***i***_) ∈ ℝ^*d*^ is the output of the encoder for the *i*-th node. Here, ***r*** ∈ ℝ^*d*^ is the mean pooled output of the encoder and the vectorized representation of the local structure.

#### Local structure codebook

The structure code book quantizes dense vectors representing protein structure into discrete tokens (Figure 1B). To build this, we employ a structure encoder to embed the local structures of all residues from the C.A.T.H dataset (See in Appendix A.2) into a continuous latent space. Then we apply the *k*-means algorithm to identify *K* centroids within this latent space, denoted as 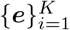 . These centroids constitute the structure codebook, as shown in Figure 1B. For any local-structure embedding, it is quantized by the nearest vector ***e***_*j*_ within the codebook and *j* serving as the structure token. In this paper, the clustering number *K* is also referred to as the structure vocabulary size.

#### Protein serialization and quantization

In general, for a residue at position *i* in a protein sequence, we first build a graph ***G***_*i*_ only based on its local structure, and then use the structure encoder to embed it into a continuous vector ***r***_*i*_. Then we use the codebook to assign a structure token *s*_*i*_ ∈ {1, 2, …, *K*} to this vector as the structure token of the residue. Overall, the entire protein structure can be serialized and quantized into a sequence of structure tokens (Figure 1C).

### 3.2 Sequence-Structure Disentangled Attention

Inspired by DeBerta [40], we use an expanded form of disentangled attention to combine the attention of residual sequences and structure sequences as well as relative positions. Specifically, for a residue at position *i* in a protein sequence, it can be represented by three items: ***R***_*i*_ denotes its residue token hidden state, ***S***_*i*_ represents the embedding of residue-level local structure, and ***P*** _*i*_|_*j*_ is the embedding of relative position with the token at position *j*. The calculation of the cross attention ***A***_*i,j*_ between residue *i* and residue *j* can be decomposed into nine components by:

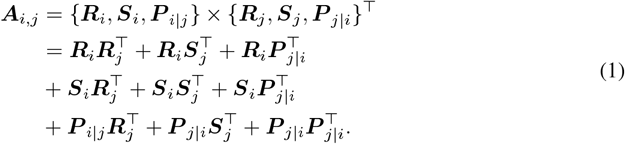

As formulated in Equation 1, the attention weight of a residue pair can be calculated by separate matrices, including residue tokens, structure tokens, and relative positions. These matrices are utilized for various interactions such as *residue-to-residue, residue-to-structure, residue-to-position, structure-to-residue, structure-to-structure, structure-to-position, position-to-residue, position-to-structure, and position-to-position*. Since our model concentrates on learning contextual embeddings for residues, the terms structure-to-structure 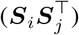, structure-to-position 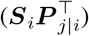, position-to-structure 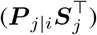, and position-to-position 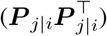 do not provide relevant information about residues and thus do not contribute significantly. Consequently, these terms are removed from our implementation of the attention weight calculation. As shown in Figure 2, our sequence-structure disentangled attention mechanism includes 5 types of attention.

**Figure 2.**
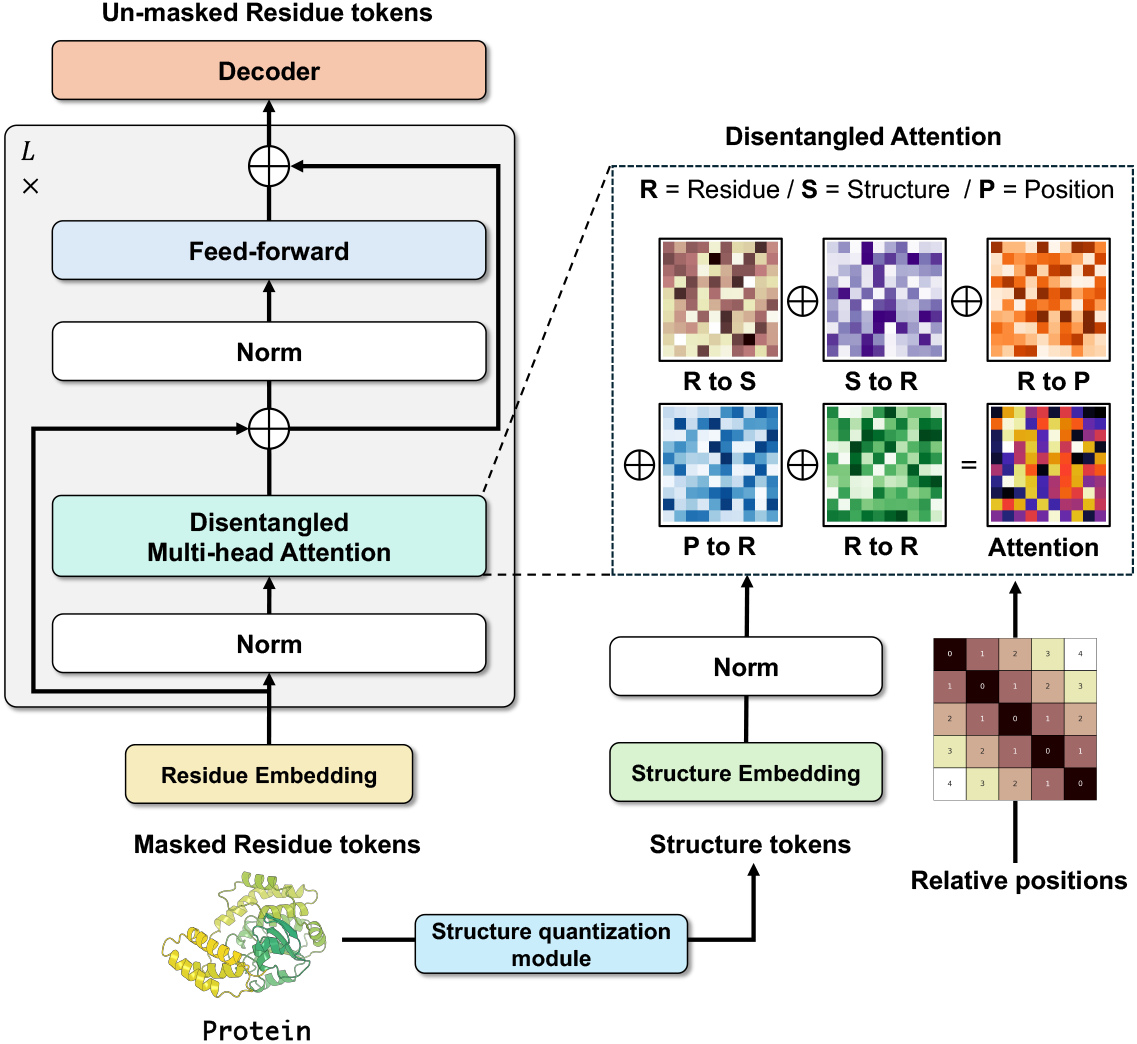
Model architecture of ProSST. ProSST is a Transformer-style model and the difference is that ProSST uses disentangled attention instead of self-attention [19].

In the following part, we use single-head attention as an example to demonstrate the operation of sequence-structure disentangled attention. To begin, we define the relative position of the *i*-th to the *j*-th residue, denoted as *δ*(*i, j*):

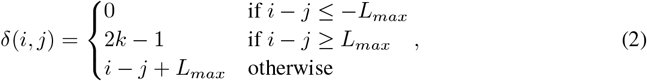

where, *L*_*max*_ represents the maximum relative distance we consider, which is set to 1024 in the implementation. Similar to standard self-attention operation [19], the computation of query, key for structure, residue and relative position, and the value for residue is as follows:

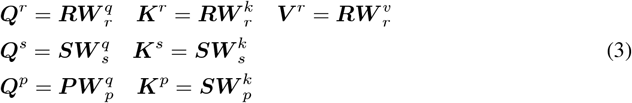

and the the attention score *Â*_*i,j*_ from residue *i* to residue *j* can be calculated as follows:

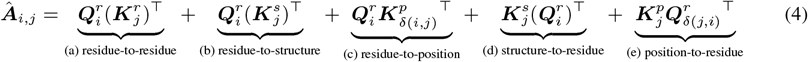

where 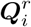 represents the *i*-th row of the matrix ***Q***^*r*^, and 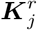 denotes the *j*-th row of ***K***^*r*^. 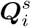 and 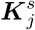 are the *i*-th and *j*-th rows of ***Q***^*s*^ and ***K***^*s*^, respectively. The term 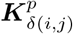 refers to the row in ***K***^*p*^ indexed by the relative distance *δ*(*i, j*), and 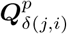 refers to the row in ***Q***^*p*^ indexed by the relative distance *δ*(*j, i*). To normalize the attention scores, a scaling factor of 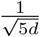 is applied to *Â* . This scaling is crucial for ensuring the stability of model training [40], particularly when dealing with large-scale language models. All the *Â*_*ij*_ form the attention matrix, and the final output residue hidden state is ***R***_*o*_:

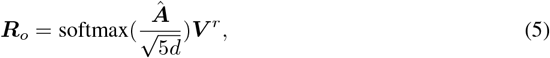

which is used as the input for the hidden state of the next layer (see Appendix 1 for the algorithm of disentangled attention).

### 3.3 Pre-Training Objective

ProSST is pre-trained with the structure-conditioned masked language modeling. In this approach, each input sequence ***x*** is noised by substituting a fraction of the residues with a special mask token or other residues. The objective of ProSST is to predict the original tokens that have been noise in the input sequence, utilizing both the corrupted sequence and its structure token sequence ***s*** as context:

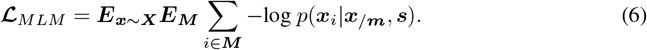

We randomly select 15% indices from the set ***M*** for nosing and computing loss for back-propagation. At each selected index *i*, there is an 80% chance of substituting the residue with a mask token, a 10% chance of replacing it with a random residue token, and the remaining residues are unchanged. The training objective is to minimize the negative log-likelihood for each noised residue ***x***_*i*_, based on the partially noised sequence ***x/M*** and the *un-noised* structure tokens, serving as contextual cues. Therefore, to accurately predict the noised tokens, this objective enables the model not only to learn the dependencies between residues but also the relationship between residues and structures. The details of pre-training dataset and hyper-parameter configuration can be found in Appendix A.2.

## 4 Experiments

In this section, we comprehensively evaluate the representation ability of ProSST in several benchmarks, covering zero-shot mutant effective prediction tasks (Section 4.1) and various supervised function prediction tasks (Section 4.2). Additionally, we also provide ablation studies and discussions to further show the effectiveness of the detailed designs in our model (Section 4.3).

### 4.1 Zero-Shot Mutant Effect Prediction

#### Datasets

To evaluate the effectiveness of ProSST in zero-shot mutant effect prediction, we conduct experiments on ProteinGym [41] and utilize AlphaFold2 [11] to generate the structures of wild-type sequences. We also evaluate ProSST on ProteinGym-Stablility, a subset of ProteinGym to assess the thermostability of proteins. See Appendix A.2 for the details of the dataset.

#### Baselines

We compare ProSST with the current state-of-the-art models, including sequence-based models [6, 7, 21], structure-sequence model [14], inverse folding models [35, 36], evolutionary models [42, 43, 44], and ensemble models [6, 45].

#### Results

Table 1 shows the performance of zero-shot mutant effect prediction on ProteinGYM. Based on the results, we draw several noteworthy conclusions:

**Table 1:**
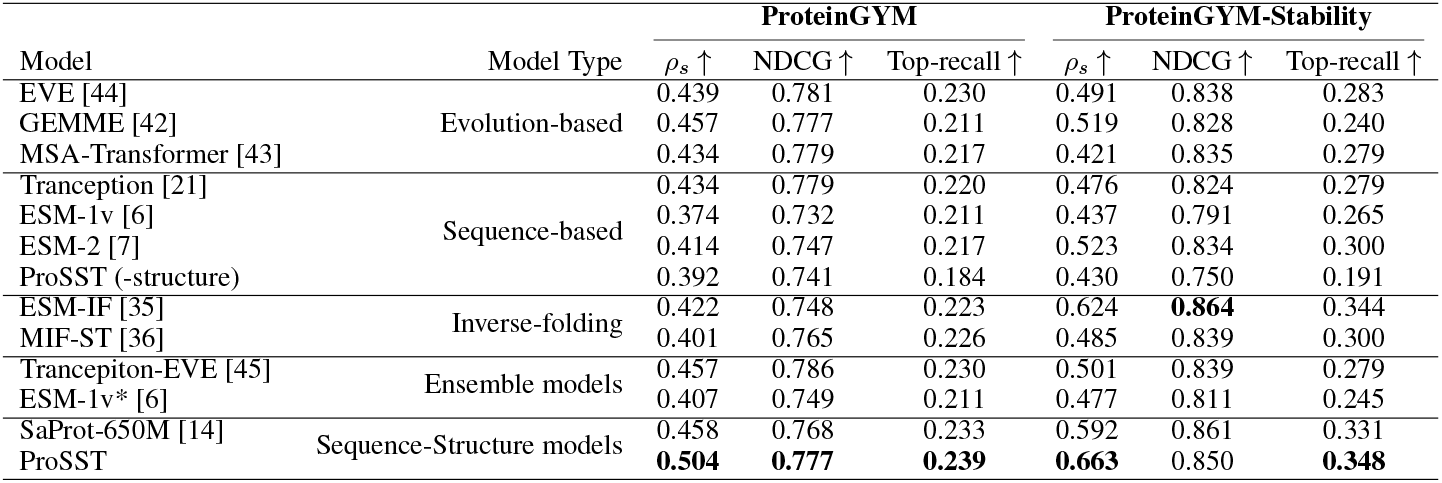
Comparison of zero-shot mutation prediction performance between ProSST and other models. *ρ*_*s*_ is the Spearman rank correlation.

- ProSST outperforms all baselines on zero-shot mutant effect predictions of ProteinGYM. Besides, it achieves the best performance in predicting stability, aligning with the previous findings that models incorporating structure information typically perform better in stability predictions [41].
- The degraded version of ProSST (without structure) gets results similar to other sequence-based models. This demonstrates that the performance improvement of our model stems from the efficient modeling of structure information, rather than other factors such as more powerful backbones.

### 4.2 Supervised Fine-Tuning Tasks

#### Downstream tasks

For supervised learning, we choose four protein downstream tasks, including thermostability prediction, Metal Ion Binding prediction, protein localization prediction (DeepLoc) and GO annotations prediction (three settings including MF, BO, and CC). More details of the tasks, datasets, and metrics can be found in Appendix A.2

#### Baselines

We compared ProSST with other PLMs including ESM-2[7], ESM-1b [5], and the sequence-structure model SaProt [14] (two parameter versions, 35M, 650M), MIF-ST [36], as well as the protein structure representation model GearNet [28].

#### Results

The results of the supervised fine-tuning tasks are shown in Table 4.2, and we can get the following conclusions:

**Table 2:** Comparison of supervised fine-tuning on downstream tasks. *ρ*_*s*_ denotes the Spearman correlation coefficient.

| Model | # Params | DeepLoc | Metal Ion Binding | Thermostability | GO-MF | GO-BP | GO-CC |
| --- | --- | --- | --- | --- | --- | --- | --- |
| | | Acc% ↑ | Acc% ↑ | $\rho_s \uparrow$ | F1-Max ↑ | F1-Max ↑ | F1-Max ↑ |
| ESM-2 | 650M | 91.96 | 71.56 | 0.680 | 0.668 | 0.345 | 0.411 |
| ESM-1b | 650M | 92.83 | 73.57 | 0.708 | 0.661 | 0.320 | 0.392 |
| MIF-ST | 643M | 91.76 | 75.08 | 0.694 | 0.627 | 0.239 | 0.248 |
| GearNet | 42M | 89.18 | 71.26 | 0.571 | 0.650 | 0.354 | 0.404 |
| SaProt-35M | 35M | 91.96 | 71.56 | 0.680 | 0.668 | 0.345 | 0.411 |
| SaProt-650M | 650M | 93.55 | 75.75 | 0.724 | **0.678** | 0.356 | 0.414 |
| ProSST | 110M | **94.68** | **76.67** | **0.724** | 0.672 | **0.387** | **0.501** |

- ProSST gets the best results among all models with 5 firsts and 1 second in all 6 settings. For the tasks (settings) of DeepLoc, Metal Ion Binding, GO-BP, and GO-CC, ProSST largely surpasses other methods, and SaProt-650M gets comparable (or slightly better) results for thermostability and GO-MF with ProSST, at the price of about 6*×* model size.
- The sequence-structure models, SaProt and ProSST, show better results than other counterparts, which suggests the importance of the structure cues in protein modeling. Furthermore, ProSST is more capable of integrating sequence and structure information of proteins than SaProt, which confirms the effectiveness of our designs.

Combined with the results in Section 4.1, ProSST exhibits powerful ability in multiple settings.

### 4.3 Ablation Study

In this section, we provide additional ablation studies and discussions to show the necessity and effectiveness of the detailed designs in ProSST. Specifically, we use zero-shot mutant effect prediction on ProteinGYM, supervised downstream task DeepLoc, and the perplexity in the pre-training validation set to conduct corresponding experiments.

#### Ablations on quantized structure

The ablation results of quantized structure are shown in Table 3 and Figure 3(a), and we can get the following findings:

- We can find, as the increases of *K* (the size of local structure vocabulary), the performance of ProSST shows an upward trend on all metrics, and most metrics achieve the best results with *K* = 2048. Based on that, we set *K* = 2048 as our default setting.
- As the increase of K, the convergence of ProSST improves progressively (Figure 3(a)), which suggests incorporating structure cues can improve the representation capabilities of models.
- Based on the same network architecture, the proposed structure quantization method (with an appropriate hyper-parameter *K*) performs better than Foldseek [16] and DSSP [37], which shows the effectiveness of our design.
- ProSST (Foldseek), ProSST (DSSP), and all ProSST (K>0) models significantly surpass ProSST (K=0) in all metrics, which confirms the importance of the structure cues again.

**Table 3:** Ablation studies on quantized structure. We first show the performance of our models with K centroids of local structures. ProSST (K=0) refers to the model without structure token sequence. We also replace the proposed quantization method with existing Foldseek and DSSP, and show the results of these variants.

|  | DeepLoc | ProteinGYM |  |  | Pretraining |
| --- | --- | --- | --- | --- | --- |
| | Acc% ↑ | $\rho_s$ ↑ | NDCG ↑ | Top-Recall ↑ | Perplexity ↓ |
| ProSST (K=4096) | 94.16 | 0.498 | 0.773 | 0.233 | **8.880** |
| **ProSST(K=2048)** | **94.68** | **0.504** | **0.777** | **0.239** | 9.033 |
| ProSST (K=1024) | 93.41 | 0.485 | 0.760 | 0.231 | 9.333 |
| ProSST (K=512) | 93.71 | 0.471 | 0.759 | 0.223 | 9.577 |
| ProSST (K=128) | 93.18 | 0.469 | 0.753 | 0.228 | 10.021 |
| ProSST (K=20) | 93.18 | 0.438 | 0.744 | 0.210 | 10.719 |
| ProSST (K=0) | 89.65 | 0.392 | 0.741 | 0.184 | 12.190 |
| ProSST (Foldseek) | 93.24 | 0.468 | 0.759 | 0.228 | 10.049 |
| ProSST (DSSP) | 93.21 | 0.439 | 0.760 | 0.204 | 10.009 |

**Figure 3.**
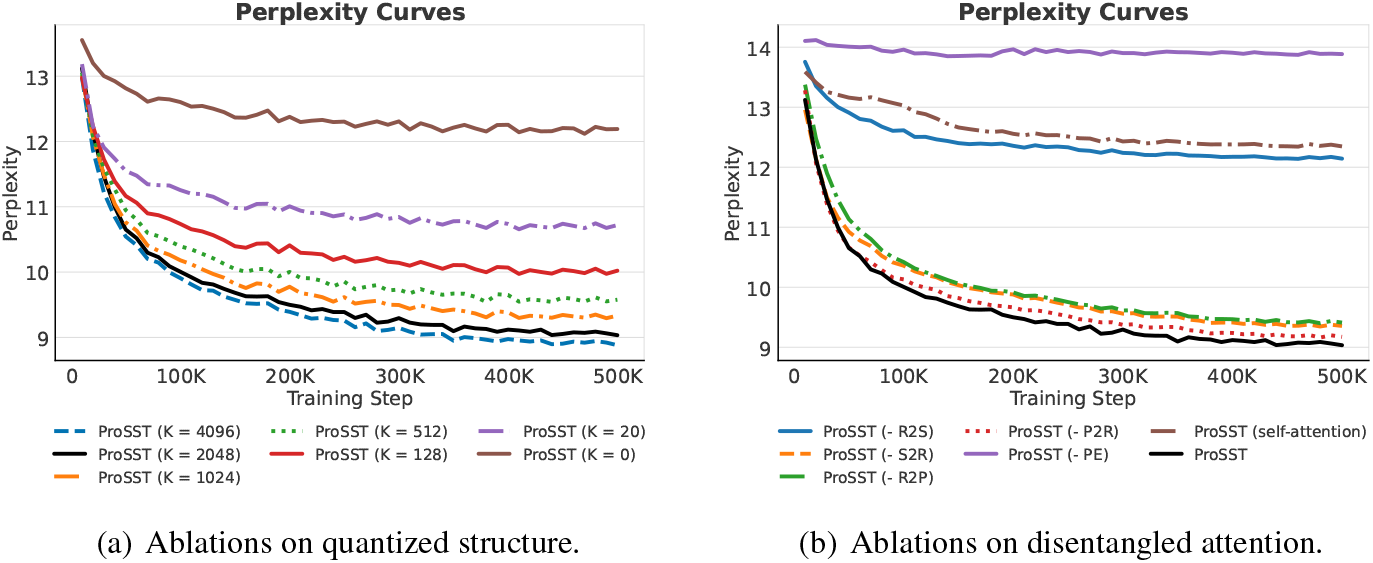
Perplexity curves of ProSST under different settings. We ablate the components of quantized structure and disentangled attention, and show their perplexity curves on the validation set.

#### Ablations on disentangled attention

Here we show detailed ablations and comparisons of disentangled attention in Table 4 and Figure 3(b), and we can get the following observations:

- All items in Equation 4 are necessary to our attention formulation. Also note that ‘P2R’ attention has the least impact on model capacity, with the Perplexity slightly increasing from 9.033 to 9.173, suggesting that positional attention to amino acids is relatively less critical than other items. Conversely, removing ‘R2S’ item results in a significant increase in Perplexity from 9.033 to 12.142, underscoring the important role of structure information in enhancing the model ‘s representation capability.
- Compared with standard self-attention, our attention formulation gets better results for all metrics, indicating that explicitly modeling structure cues is crucial for integrating such information. Besides, positional encoding is also necessary in our design.

**Table 4:** Ablation studies on disentangled attention. The term “-S2R” denotes the removal of structure-to-residue in our attention formulation, similar to other terms, and “-PE” denotes the removal of positional encoding. ProSST (self-attention) refers to the model trained with standard attention (with structure cues).

|  | DeepLoc | ProteinGYM |  |  | Pretraining |
| --- | --- | --- | --- | --- | --- |
| | Acc% ↑ | $\rho_s$ ↑ | NDCG ↑ | Top-Recall ↑ | Perplexity ↓ |
| ProSST | **94.68** | **0.502** | **0.793** | **0.252** | **9.033** |
| ProSST (- P2R) | 91.46 | 0.478 | 0.778 | 0.227 | 9.173 |
| ProSST (- R2P) | 92.46 | 0.466 | 0.772 | 0.216 | 9.410 |
| ProSST (- R2S) | 92.19 | 0.438 | 0.766 | 0.208 | 12.142 |
| ProSST (- S2R) | 91.38 | 0.475 | 0.779 | 0.226 | 9.355 |
| ProSST (- PE) | 86.06 | 0.095 | 0.634 | 0.126 | 13.885 |
| ProSST (self-attention) | 90.42 | 0.401 | 0.728 | 0.189 | 12.346 |

## 5 Conclusion and Limitations

This paper introduces ProSST, a protein sequence-structure transformer for PLM. ProSST includes two key techniques, protein structure quantization module and sequence-structure disentangled attention. The structure quantization module contains an encoder and a k-means clustering model. The encoder is trained with a denoising objective and is utilized for encoding protein structures. Leveraging this encoder, we embed the local structures of each residue within every protein in the C.A.T.H dataset into a continuous latent space. Then we utilize k-means clustering algorithm to obtain *K* (default setting is 2048) centroids. These centroids are then utilized to discretize the local structures of residues based on the index of the nearest centroid of its structure embedding vectors.

A protein structure can be transformed into a sequence of discrete numbers (or referred to tokens) and each token representing the corresponding local structure of residue. The sequence-structure attention enhances standard self-attention by not only considering self-attention residues but also incorporating attention between residues and structures, and vice versa. This enables the model to learn the relationships between residues and structures, thereby acquiring improved adequate contextual representations of residues. Furthermore, we pre-train ProSST with 18.8 million protein structures using a MLM objective. Experimental results show that ProSST can outperform existing models in ProteinGYM benchmark and other supervised learning tasks. Despite of this, there are some limitations of ProSST. For example, the local structure construction and encoding requires heavy computations. In the future work, we aim to speed up the protein structure quantization process. Additionally, we plan to enhance ProSST by training it with larger structure datasets and expanding its parameter, which may further improve its performance.

## 6 Acknowledgements

This work was supported by the grants from the National Science Foundation of China (grant number 12104295), the Computational Biology Program of Shanghai Science and Technology Commission (23JS1400600), Shanghai Jiao Tong University Scientific and Technological Innovation Funds (21X010200843), the Student Innovation Center at Shanghai Jiao Tong University, and Shanghai Artificial Intelligence Laboratory

## A. Appendix

### A.1 Zero-Shot Scoring

Previous studies have demonstrated that PLMs, when trained on extensive and varied protein sequence databases, are capable of predicting experimental measurements of protein mutants function without further supervision. For those PLMs that are trained with masked language modeling objective, the calculation of mutation scores can be formalized as follows:

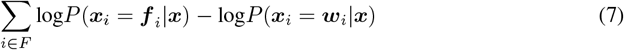

Here ***F*** is a mutant with multi-point the mutation, and ***F*** = {(*i*, ***F*** _*i*_, ***w***_*i*_) |*i* = 1, 2, …, |***F***|} is a set of triplets, where *i* ∈ℕ represents the position of a point mutation, ***F*** _*i*_ is the residue after the point mutation, and ***w***_*i*_ is the original residue of the point mutation. ***x*** is the sequence of amino acids of the wild type. We slightly modify the formula above to adapt to ProSST, where the structure sequence is an additional condition to score mutants:

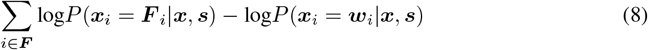

Here, ***s*** is the structure token sequence of the wild type.

### A.2 Details of the Datasets and Metrics

#### Dataset for pre-training

The pre-training data is collected from AlphaFoldDB [13], which contains more than 214 million structures predicted by AlphaFold [11]. We downloaded the 90% reduced version, containing 18.8 million structures.^3^. From this collection, we randomly select 100,000 structures for validation, enabling us to monitor the perplexity in the training phase.

#### Dataset for training structure encoder

The dataset used for training the structure encoder originates from CATH43-S40 ^4^. This dataset is manually annotated and comprises protein crystal structural domains that have been deduplicated for sequence similarity by 40%. The original dataset contains 31,885 structures. After removing structural domains missing atoms such as ***C*α** and N, the dataset is reduced to 31,270 entries. From this, 200 structures were randomly selected to serve as a validation set. The auto-encoder model was then trained using the configuration that yielded the lowest loss on this validation set.

#### Dataset for training structure codebook

The dataset for training the structure codebook consists of local structures extracted from CATH43-S40. Given a protein structure, slide along the residue sequence to select a segment with a chosen residue as the anchor. Connect up to 40 residues within 10 Å to form a star-shaped graph. For pairwise amino acid pairs in this graph, if the Euclidean distance is less than 10 Å, a link will be assigned to them. This process yields a number of protein local structures equal to the length of the protein multiplied by the total number of proteins, resulting in 4,735,677 local structures from the protein structures in CATH43-S40. These sub-structures are fed into a structural encoder to obtain embeddings. By setting various quantities for *K*, different structure codebooks are obtained using the k-means clustering algorithm.

#### Dataset and metrics for zero-shot mutant effect prediction

We utilize the ProteinGYM benchmark [41] to assess the zero-shot mutant effect prediction capabilities of ProSST. ProteinGYM offers a comprehensive benchmarks specifically collected for predicting protein fitness. It contains a wide range of deep mutational scanning assays with millions of mutated sequences. ProSST is evaluated using the most extensively utilized datasets for substitution mutations, which include 217 experimental assays. Each assay incorporates both the sequence and structure of the protein, with a particular emphasis on 66 datasets that focus on thermo-stability. The evaluation metrics employed are the Spearman coefficient, Top-recall, and NDCG, where higher values signify superior model performance. These metrics are computed using scripts ^5^ provided by ProteinGYM.

#### Datasets and metrics for downstream tasks

- **Thermostability**. The task is to predict the thermostability values of proteins using the “Human-cell” divisions from the Thermostability task in FLIP [46]. For this regression task, the Spearman correlation coefficient is utilized as the evaluation metric to evaluate the prediction results.
- **DeepLoc** (Protein Sub-cellular Localization). The task is to output a probability distribution across two sub-cellular localization categories for a protein. This is a binary classification task, and we utilize accuracy as the metric to evaluate the predictions. This dataset was introduced by DeepLoc [47] and we use the original data split.
- **Metal Ion Binding**. The task is to predict whether there exist metal ion-binding sites within a protein. This is also a binary classification task, and we utilize accuracy as the metric to evaluate the predictions. This dataset was introduced by TAPE [20], and we use the original data split.
- **GO annotations prediction**. This task is to predict Gene Ontology terms to evaluate the model ‘s ability to predict protein functions. This task was introduced by DeepFRI [26], and we use three types of GO labels: Molecular Function (MF), Biological Process (BP), and Cellular Component (CC). This is a multi-label classification task, and we evaluate the model using the Max F1-Score.

### A.3 Details of Implementations

#### Structure encoder

We describe a structure with the graph ***G*** = (***V***, ***E***), adopting the characterizations of ***V*** and ***E*** as outlined in the GVP framework [18], excluding the one-hot representation of residue. The dimensions for node and edge representations are set at 256 and 64, respectively, with the encoder comprising six layers. For optimization, we employ the Adam optimizer in a mini-batch gradient descent approach. To manage computational load, batches are formed by grouping structures of similar sizes, with each batch containing no more than 3000 nodes. The learning rate is set to 10^−4^. The dropout probability is set to 10^−4^. And The number of graph layers is set at 6.

#### Pre-training

All ProSST models is trained on a DGX-A800 GPU (8×80G) server in BF16 precision for about a month. The model has 12 transformer layers, 12 attention heads, and 768 embedding dims with 3172 feed-forward embedding dimensions with the GELU activation function. We train with 4096 tokens per mini-batch for 500,000 steps. We use AdamW [48] as our optimizer with *β*_1_ and *β*_2_ set to 0.9 and 0.999, and a weight decay value of 0.001. We warm up the learning rate from 0 to 0.0002 over the first 2000 steps, then decay it by a cosine schedule to the 0. We use a dropout rate of 0.1 and clip gradients using a clipping value of 1.0. For the tokenization of the protein data, we use the residue-level tokenizer which is adopted in several PLMs [5, 7, 6]. To make the structure sequence the same length as the amino acid sequence, we also added special *[SOS], [EOS]*, and *[PAD]* token for the structure sequences.

#### Fine-tuning

To ensure fair comparisons, we fine-tuned ProSST using a fixed set of hyper-parameters. We use for the Adam optimizer with *β*_1_ set to 0.9, *β*_2_ to 0.98, and applied an *L*2 weight decay of 0.001. The batch size was maintained at 64 (If 64 causes the GPU memory to explode, we will reduce the batch size and then use gradient accumulation to achieve the same batch size.) and the learning rate was set at 0.00003, except for Go annotation prediction, where it was adjusted to 0.00001. We fine-tuned all model parameters for 200 epochs, and we choose the best checkpoints based on validation set performance. Following SaProt [14]^6^, we downloaded all protein structures identified by Uniprot IDs from AFDB [13], and any proteins not found in AFDB were excluded.

### A.4 Sequence-Structure Disentangled Attention Algorithm

Here we detail the algorithm for structure-sequence disentangled attention in Algorithm 1.

#### Algorithm 1

Sequence-Structure Disentangled Attention Algorithm

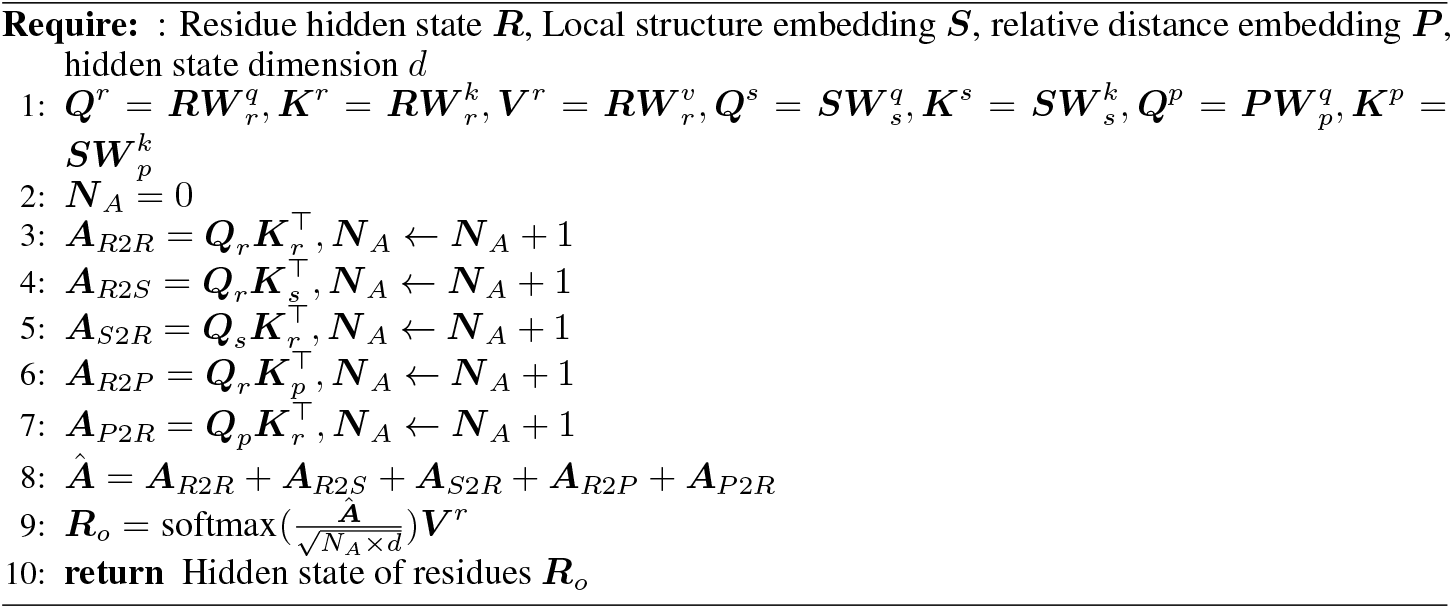

## Footnotes

2 https://github.com/ai4protein/ProSST

3 https://cluster.foldseek.com/

4 http://download.cathdb.info/cath/releases/all-releases/v4_3_0/non-redundant-data-sets/

5 https://github.com/OATML-Markslab/ProteinGym/blob/main/scripts/

6 https://github.com/westlake-repl/SaProt

## References

[1] William P Jencks. Catalysis in chemistry and enzymology. Courier Corporation, 1987.

[2] The UniProt Consortium. UniProt: the Universal Protein Knowledgebase in 2023. Nucleic Acids Research, 51(D1):D523–D531, 11 2022.

[3] Alec Radford, Jeffrey Wu, Rewon Child, David Luan, Dario Amodei, Ilya Sutskever, et al. Language models are unsupervised multitask learners. OpenAI blog, 1(8):9, 2019.

[4] Jacob Devlin, Ming-Wei Chang, Kenton Lee, and Kristina Toutanova. Bert: Pre-training of deep bidirectional transformers for language understanding. arXiv preprint arXiv:1810.04805, 2018.

[5] Alexander Rives, Joshua Meier, Tom Sercu, Siddharth Goyal, Zeming Lin, Jason Liu, Demi Guo, Myle Ott, C. Lawrence Zitnick, Jerry Ma, and Rob Fergus. Biological structure and function emerge from scaling unsupervised learning to 250 million protein sequences. Proceedings of the National Academy of Sciences, 118(15):e2016239118, 2021.

[6] Joshua Meier, Roshan Rao, Robert Verkuil, Jason Liu, Tom Sercu, and Alex Rives. Language models enable zero-shot prediction of the effects of mutations on protein function. In M. Ranzato, A. Beygelzimer, Y. Dauphin, P.S. Liang, and J. Wortman Vaughan, editors, Advances in Neural Information Processing Systems,volume 34, pages 29287–29303. Curran Associates, Inc., 2021.

[7] Zeming Lin, Halil Akin, Roshan Rao, Brian Hie, Zhongkai Zhu, Wenting Lu, Nikita Smetanin, Robert Verkuil, Ori Kabeli, Yaniv Shmueli, et al. Evolutionary-scale prediction of atomic-level protein structure with a language model. Science, 379(6637):1123–1130, 2023.

[8] Ahmed Elnaggar, Michael Heinzinger, Christian Dallago, Ghalia Rehawi, Yu Wang, Llion Jones, Tom Gibbs, Tamas Feher, Christoph Angerer, Martin Steinegger, et al. Prottrans: Toward understanding the language of life through self-supervised learning. IEEE transactions on pattern analysis and machine intelligence, 44(10):7112–7127, 2021.

[9] Nadav Brandes, Dan Ofer, Yam Peleg, Nadav Rappoport, and Michal Linial. Proteinbert: a universal deep-learning model of protein sequence and function. Bioinformatics, 38(8):2102– 2110, 2022.

[10] Hedi Hegyi and Mark Gerstein. The relationship between protein structure and function: a comprehensive survey with application to the yeast genome 11edited by g. von heijne. Journal of Molecular Biology, 288(1):147–164, 1999.

[11] John Jumper, Richard Evans, Alexander Pritzel, Tim Green, Michael Figurnov, Olaf Ronneberger, Kathryn Tunyasuvunakool, Russ Bates, Augustin Žídek, Anna Potapenko, et al. Highly accurate protein structure prediction with alphafold. Nature, 596(7873):583–589, 2021.

[12] Minkyung Baek, Frank DiMaio, Ivan Anishchenko, Justas Dauparas, Sergey Ovchinnikov, Gyu Rie Lee, Jue Wang, Qian Cong, Lisa N Kinch, R Dustin Schaeffer, et al. Accurate prediction of protein structures and interactions using a three-track neural network. Science, 373(6557):871–876, 2021.

[13] Mihaly Varadi, Stephen Anyango, Mandar Deshpande, Sreenath Nair, Cindy Natassia, Galabina Yordanova, David Yuan, Oana Stroe, Gemma Wood, Agata Laydon, et al. Alphafold protein structure database: massively expanding the structural coverage of protein-sequence space with high-accuracy models. Nucleic acids research, 50(D1):D439–D444, 2022.

[14] Jin Su, Chenchen Han, Yuyang Zhou, Junjie Shan, Xibin Zhou, and Fajie Yuan. Saprot: Protein language modeling with structure-aware vocabulary. In The Twelfth International Conference on Learning Representations, 2024.

[15] Michael Heinzinger, Konstantin Weissenow, Joaquin Gomez Sanchez, Adrian Henkel, Martin Steinegger, and Burkhard Rost. Prostt5: Bilingual language model for protein sequence and structure. bioRxiv, pages 2023–07, 2023.

[16] Michel Van Kempen, Stephanie S Kim, Charlotte Tumescheit, Milot Mirdita, Jeongjae Lee, Cameron LM Gilchrist, Johannes Söding, and Martin Steinegger. Fast and accurate protein structure search with foldseek. Nature Biotechnology, pages 1–4, 2023.

[17] Hongyuan Lu, Daniel J Diaz, Natalie J Czarnecki, Congzhi Zhu, Wantae Kim, Raghav Shroff, Daniel J Acosta, Bradley R Alexander, Hannah O Cole, Yan Zhang, et al. Machine learning-aided engineering of hydrolases for pet depolymerization. Nature, 604(7907):662–667, 2022.

[18] Bowen Jing, Stephan Eismann, Patricia Suriana, Raphael John Lamarre Townshend, and Ron Dror. Learning from protein structure with geometric vector perceptrons. In International Conference on Learning Representations, 2021.

[19] Ashish Vaswani, Noam Shazeer, Niki Parmar, Jakob Uszkoreit, Llion Jones, Aidan N Gomez, Łukasz Kaiser, and Illia Polosukhin. Attention is all you need. In I. Guyon, U. Von Luxburg, S. Bengio, H. Wallach, R. Fergus, S. Vishwanathan, and R. Garnett, editors, Advances in Neural Information Processing Systems, volume 30. Curran Associates, Inc., 2017.

[20] Roshan Rao, Nicholas Bhattacharya, Neil Thomas, Yan Duan, Peter Chen, John Canny, Pieter Abbeel, and Yun Song. Evaluating protein transfer learning with tape. In H. Wallach, H. Larochelle, A. Beygelzimer, F. d’Alché-Buc, E. Fox, and R. Garnett, editors, Advances in Neural Information Processing Systems, volume 32. Curran Associates, Inc., 2019.

[21] Pascal Notin, Mafalda Dias, Jonathan Frazer, Javier Marchena Hurtado, Aidan N Gomez, Debora Marks, and Yarin Gal. Tranception: protein fitness prediction with autoregressive transformers and inference-time retrieval. In International Conference on Machine Learning, pages 16990–17017. PMLR, 2022.

[22] Erik Nijkamp, Jeffrey A Ruffolo, Eli N Weinstein, Nikhil Naik, and Ali Madani. Progen2: exploring the boundaries of protein language models. Cell systems, 14(11):968–978, 2023.

[23] Noelia Ferruz, Steffen Schmidt, and Birte Höcker. Protgpt2 is a deep unsupervised language model for protein design. Nature communications, 13(1):4348, 2022.

[24] Ahmed Elnaggar, Hazem Essam, Wafaa Salah-Eldin, Walid Moustafa, Mohamed Elkerdawy, Charlotte Rochereau, and Burkhard Rost. Ankh: Optimized protein language model unlocks general-purpose modelling. arXiv preprint arXiv:2301.06568, 2023.

[25] Bo Chen, Xingyi Cheng, Pan Li, Yangli-ao Geng, Jing Gong, Shen Li, Zhilei Bei, Xu Tan, Boyan Wang, Xin Zeng, et al. xtrimopglm: unified 100b-scale pre-trained transformer for deciphering the language of protein. arXiv preprint arXiv:2401.06199, 2024.

[26] Vladimir Gligorijević, P Douglas Renfrew, Tomasz Kosciolek, Julia Koehler Leman, Daniel Berenberg, Tommi Vatanen, Chris Chandler, Bryn C Taylor, Ian M Fisk, Hera Vlamakis, et al. Structure-based protein function prediction using graph convolutional networks. Nature communications, 12(1):3168, 2021.

[27] Pedro Hermosilla, Marco Schäfer, Matej Lang, Gloria Fackelmann, Pere-Pau Vázquez, Barbora Kozlikova, Michael Krone, Tobias Ritschel, and Timo Ropinski. Intrinsic-extrinsic convolution and pooling for learning on 3d protein structures. In International Conference on Learning Representations, 2021.

[28] Zuobai Zhang, Minghao Xu, Arian Rokkum Jamasb, Vijil Chenthamarakshan, Aurelie Lozano, Payel Das, and Jian Tang. Protein representation learning by geometric structure pretraining. In The Eleventh International Conference on Learning Representations, 2023.

[29] Bingxin Zhou, Lirong Zheng, Banghao Wu, Yang Tan, Outongyi Lv, Kai Yi, Guisheng Fan, and Liang Hong. Protein engineering with lightweight graph denoising neural networks. bioRxiv, pages 2023–11, 2023.

[30] Ruidong Wu, Fan Ding, Rui Wang, Rui Shen, Xiwen Zhang, Shitong Luo, Chenpeng Su, Zuofan Wu, Qi Xie, Bonnie Berger, et al. High-resolution de novo structure prediction from primary sequence. BioRxiv, pages 2022–07, 2022.

[31] Zichen Wang, Steven A Combs, Ryan Brand, Miguel Romero Calvo, Panpan Xu, George Price, Nataliya Golovach, Emmanuel O Salawu, Colby J Wise, Sri Priya Ponnapalli, et al. Lm-gvp: an extensible sequence and structure informed deep learning framework for protein property prediction. Scientific reports, 12(1):6832, 2022.

[32] Zuobai Zhang, Minghao Xu, Aurelie Lozano, Vijil Chenthamarakshan, Payel Das, and Jian Tang. Enhancing protein language model with structure-based encoder and pre-training. In ICLR 2023 - Machine Learning for Drug Discovery workshop, 2023.

[33] Yang Tan, Bingxin Zhou, Lirong Zheng, Guisheng Fan, and Liang Hong. Semantical and topological protein encoding toward enhanced bioactivity and thermostability. bioRxiv, pages 2023–12, 2023.

[34] Victor Garcia Satorras, Emiel Hoogeboom, and Max Welling. E (n) equivariant graph neural networks. In International conference on machine learning, pages 9323–9332. PMLR, 2021.

[35] Chloe Hsu, Robert Verkuil, Jason Liu, Zeming Lin, Brian Hie, Tom Sercu, Adam Lerer, and Alexander Rives. Learning inverse folding from millions of predicted structures. In International conference on machine learning, pages 8946–8970. PMLR, 2022.

[36] Kevin K Yang, Niccolò Zanichelli, and Hugh Yeh. Masked inverse folding with sequence transfer for protein representation learning. Protein Engineering, Design and Selection, 36:gzad015, 2023.

[37] Wolfgang Kabsch and Christian Sander. Dictionary of protein secondary structure: pattern recognition of hydrogen-bonded and geometrical features. Biopolymers: Original Research on Biomolecules, 22(12):2577–2637, 1983.

[38] Joseph L Watson, David Juergens, Nathaniel R Bennett, Brian L Trippe, Jason Yim, Helen E Eisenach, Woody Ahern, Andrew J Borst, Robert J Ragotte, Lukas F Milles, et al. De novo design of protein structure and function with rfdiffusion. Nature, 620(7976):1089–1100, 2023.

[39] Ian Sillitoe, Nicola Bordin, Natalie Dawson, Vaishali P Waman, Paul Ashford, Harry M Scholes, Camilla SM Pang, Laurel Woodridge, Clemens Rauer, Neeladri Sen, et al. Cath: increased structural coverage of functional space. Nucleic acids research, 49(D1):D266–D273, 2021.

[40] Pengcheng He, Xiaodong Liu, Jianfeng Gao, and Weizhu Chen. Deberta: Decoding-enhanced bert with disentangled attention. In International Conference on Learning Representations, 2021.

[41] Pascal Notin, Aaron Kollasch, Daniel Ritter, Lood van Niekerk, Steffanie Paul, Han Spinner, Nathan Rollins, Ada Shaw, Rose Orenbuch, Ruben Weitzman, Jonathan Frazer, Mafalda Dias, Dinko Franceschi, Yarin Gal, and Debora Marks. Proteingym: Large-scale benchmarks for protein fitness prediction and design. In A. Oh, T. Neumann, A. Globerson, K. Saenko, M. Hardt, and S. Levine, editors, Advances in Neural Information Processing Systems, volume 36, pages 64331–64379. Curran Associates, Inc., 2023.

[42] Elodie Laine, Yasaman Karami, and Alessandra Carbone. Gemme: a simple and fast global epistatic model predicting mutational effects. Molecular biology and evolution, 36(11):2604– 2619, 2019.

[43] Roshan M Rao, Jason Liu, Robert Verkuil, Joshua Meier, John Canny, Pieter Abbeel, Tom Sercu, and Alexander Rives. Msa transformer. In International Conference on Machine Learning, pages 8844–8856. PMLR, 2021.

[44] Jonathan Frazer, Pascal Notin, Mafalda Dias, Aidan Gomez, Joseph K Min, Kelly Brock, Yarin Gal, and Debora S Marks. Disease variant prediction with deep generative models of evolutionary data. Nature, 599(7883):91–95, 2021.

[45] Pascal Notin, Lood Van Niekerk, Aaron W Kollasch, Daniel Ritter, Yarin Gal, and Debora Susan Marks. TranceptEVE: Combining family-specific and family-agnostic models of protein sequences for improved fitness prediction. In NeurIPS 2022 Workshop on Learning Meaningful Representations of Life, 2022.

[46] Christian Dallago, Jody Mou, Kadina E Johnston, Bruce Wittmann, Nick Bhattacharya, Samuel Goldman, Ali Madani, and Kevin K Yang. FLIP: Benchmark tasks in fitness landscape inference for proteins. In Thirty-fifth Conference on Neural Information Processing Systems Datasets and Benchmarks Track (Round 2), 2021.

[47] José Juan Almagro Armenteros, Casper Kaae Sønderby, Søren Kaae Sønderby, Henrik Nielsen, and Ole Winther. Deeploc: prediction of protein subcellular localization using deep learning. Bioinformatics, 33(21):3387–3395, 2017.

[48] Ilya Loshchilov and Frank Hutter. Decoupled weight decay regularization. In International Conference on Learning Representations, 2019.

